# Dynamic formation of a posterior-to-anterior peak-alpha-frequency gradient driven by two distinct processes

**DOI:** 10.1101/2024.01.31.578276

**Authors:** Max Kailler Smith, Marcia Grabowekcy, Satoru Suzuki

## Abstract

Peak-alpha frequency varies across individuals and mental states, but it also forms a negative gradient from posterior to anterior regions in association with increases in cortical thickness and connectivity reflecting the cortical hierarchy in temporal integration. Tracking the spatial standard deviation of peak-alpha frequency in scalp EEG, we observed that the posterior-to-anterior gradient dynamically formed and dissolved. Periods of high spatial standard deviation yielded robustly negative posterior-to-anterior gradients—the gradient state—while periods of low spatial standard deviation yielded globally converged peak-alpha frequency—the uniform state. Our analyses suggest that the fluctuations between the gradient and uniform states are associated with two separate variables: (1) coordinated variations in peak-alpha frequency in anterior regions and (2) coordinated variations in peak-alpha power in central regions driven by posterior regions. These variables each accounted for ∼25% of the state fluctuations, were uncorrelated with each other, and together accounted for ∼50% of the state fluctuations. These results reflect general mechanisms as they replicated while participants engaged in a variety of behavioral tasks with their eyes closed (breath focus, vigilance, working memory, mental arithmetic, and generative thinking). Overall, our results suggest that the spatial pattern of peak-alpha frequency dynamically fluctuates between the gradient state, potentially facilitating information influx and temporal integration toward anterior regions, and the uniform state, potentially facilitating global communication, with the state fluctuations controlled by at least two distinct mechanisms, an anterior mechanism that directly adjusts peak-alpha frequencies and a posterior mechanism that indirectly adjusts them by influencing synchronization.

## Introduction

Alpha-band oscillations are the most salient and ubiquitous features of oscillatory neural activity detected in non-invasive recordings of electrophysiological activity (e.g., EEG, MEG) in humans. Accordingly, peak-alpha frequency, that is, the frequency at which oscillatory power is maximum within the alpha range, has long been a focus of human electrophysiological research. Numerous studies have demonstrated that peak-alpha frequency stably differs across individuals, increases in childhood but decreases after ∼20 years of age, tends to increase with task demands and arousal but decreases with continued task engagement, varies in relation to emotional valence, and influences sensory processing (see [1] for a review up to 2017; [2-4] for some recent results).

While a large body of research has elucidated the trait, state, and modulatory aspects of peak-alpha frequency, recent studies demonstrated a link between the spatial distribution of peak-alpha frequency and general cortical gradients of neuroanatomical and neurophysiological features. Specifically, a recent MEG study demonstrated that peak-alpha frequency generally forms a negative gradient from posterior to anterior regions, slowing toward anterior regions in association with increases in cortical thickness [5]. Increases in cortical thickness have been linked to increased connectivity [6] as well as increased ratio of feedback to feedforward connections [7], which may contribute to increased temporal integration [8-12]. These associations suggested that the negative posterior-to-anterior gradient of peak-alpha frequency reflects the cortical hierarchy in temporal integration [5,13]. Further, the gradient may facilitate information flows from posterior to anterior regions given that the negative posterior-to-anterior gradient is likely accompanied by traveling waves that follow the gradient [14-15].

In the current study, we consistently replicated the negative posterior-to-anterior gradient of peak-alpha frequency in scalp EEG while participants engaged in a variety of behavioral tasks with their eyes closed (usually generating minimal muscle artifacts), including breath focus, vigilance, working memory, mental arithmetic, and generative thinking. Interestingly, we observed that the gradient dynamically formed and dissolved. Tracking the spatial variability (measured as the spatial standard deviation) of peak-alpha frequency with reasonable temporal- (∼370 ms) and spectral-resolution (∼1 Hz), we observed that the spatial distribution of peak-alpha frequency fluctuated between a *uniform state* where peak-alpha frequency values globally converged at ∼11 Hz, and a *gradient state* where they gradually decreased from ∼11 Hz in posterior regions to ∼8 Hz in anterior regions. This fluctuation demonstrates that, while the negative posterior-to-anterior gradient, potentially reflective of the cortical hierarchy in temporal integration, is consistently observed in time-averaged EEG and MEG activity, there are also mechanisms that dynamically form and dissolve the gradient (anticipated by [5,13]).

The goal of the current study was to identify the processes that drive these state fluctuations. By applying various sensible spatial filters (including those derived from principal components), we discovered that the state fluctuations are independently associated with (1) coordinated variations in peak-alpha frequency in anterior regions and (2) coordinated variations in peak-alpha power in central regions driven by posterior regions, with the two variables accounting for as much as ∼50% of the variance in the state fluctuations. Overall, our results suggest that the spatial pattern of peak-alpha frequency dynamically fluctuates between the gradient state, potentially facilitating information influx and temporal integration toward anterior regions, and the uniform state, potentially facilitating global communication, with the state fluctuations controlled by at least two distinct mechanisms, an anterior mechanism that directly adjusts peak-alpha frequency values and a posterior mechanism that indirectly adjusts them by influencing synchronization.

## Materials and methods

### Participants and behavioral tasks

Thirty Northwestern University students and individuals from the Evanston/Chicagoland community (20 females; ages 18-39 years, *M*=23.9, *SD*=5.3) participated. They were tested individually in a dimly lit room during the period between 1/17/2019 and 7/20/2021. The study was approved by the Northwestern University Institutional Review Board and each participant provided written consent. Data from one participant was excluded from analyses because they unpredictably burst into laughter during the experiment, so our sample size was 29. Each participant performed five cognitive tasks in a fixed order with their eyes closed with pink noise playing through speakers located on a table in front of the participant. Each task was performed continuously for three minutes. In the Breath-focus task, participants were instructed to focus on the most salient physical sensations of their breath while maintaining natural breathing. In the Vigilance task, participants were instructed to listen to background pink noise to detect 15 instances (whose intervals were pseudo-randomly varied between 4 sec and 20 sec) of a brief (200 ms) volume decrement (by 15%) by pressing a button upon detection of each volume decrement. In the 2-Back task, an auditory letter was presented every 2 seconds and participants were instructed to count the number of instances of a letter being the same as the one presented 2 letters prior. In the Countdown-by-7 task, participants were instructed to count down from a given four-digit number by sevens and to report the final number at the end of the three-minute period. In the Alternative-use task, participants were instructed to think about unusual uses of a common object (e.g., shoe) throughout the three-minute period and to report what they thought was the most unusual use at the end. These tasks are not described in greater detail as the purpose of including them was to demonstrate the task invariance (generalizability) of our EEG results. Importantly, we chose a set of tasks that engaged a variety of cognitive processes including breath focus, vigilance, working memory, mental arithmetic, and generative thinking.

### EEG recording and pre-processing

While participants engaged in the Breath-focus, Vigilance, 2-Back, Countdown-by-7, and Alternative-use tasks with their eyes closed, EEG was recorded from 64 scalp electrodes (although we used a 64-electrode montage, we excluded signals from noise-prone electrodes, *Fpz, Iz, T9*, and *T10* from analyses) at a sampling rate of 512 Hz using a BioSemi ActiveTwo system (see www.biosemi.com for details). Electrooculographic (EOG) activity was monitored using four face electrodes, one placed lateral to each eye and one placed beneath each eye. Two additional reference electrodes were placed on the left and right mastoids. The EEG data were preprocessed using EEGLAB and ERPLAB toolboxes for MATLAB [16,17]. They were re-referenced offline to the average of the two mastoid electrodes, bandpass-filtered at 0.01 Hz-80 Hz, and notch-filtered at 60 Hz (to remove power-line noise that affected the EEG signals from some participants). As EEG signals were recorded with eyes closed, they were relatively free of muscle artifact. Nevertheless, we visually inspected the EEG waveforms (after applying the surface-Laplacian transform and taking the temporal derivative; see below) and the time series of peak-alpha power from all sites per condition per participant, and removed intervals that appeared to contain EEG artifacts that affected peak-alpha power. The cleanup resulted in the removal of 0.6% of the data on average as well as the interpolation of one noisy electrode, POz, by replacing its signals with those averaged from its neighbors, Pz, PO3, PO4, and Oz, for one participant in the Countdown-by-7 task. We note that the pattern of results was virtually identical with or without removing these suspected EEG artifacts.

### Estimating dura sources by surface-Laplacian transforming EEG signals

EEG source reconstruction methods constrained by structural MRI and fMRI localizers obtained from each participant may achieve superior source reconstruction with models customized for each participant [18]. Such an approach, however, was unavailable to us as we had neither structural MRI nor fMRI data for our participants. Among the non-customized source-imaging methods, we chose the surface-Laplacian transform that (theoretically) estimates the spatial distribution of macroscopic current sources/sinks on the dura surface. The surface-Laplacian transform has been shown to produce similar dura sources to those inferred by deconvolving scalp EEG potentials using a generic model of thicknesses and impedances of scalp and skull [19]. Popular source-imaging methods such as sLORETA and Beamforming have been shown to approximate simulated sources and/or to extract neural correlates of behaviors to a similar degree as the surface-Laplacian transform [20,21]. Further, there is no evidence (to our knowledge) to suggest that these popular source-imaging methods provide greater spatial resolution than the surface-Laplacian transform. Thus, our preference was to use the latter because it is the most general source-imaging method that relies the least on model-specific assumptions and free parameters [19,20,22-24].

The surface-Laplacian transform is expected to reduce volume-conduction effects from substantially greater than 5 cm in raw EEG to within 1-3 cm [19,20,25,31] which approximately corresponded to the average spacing of electrodes in our 64-channel montage. For our implementation of the surface-Laplacian transform, we used Perrin’s and colleague’s algorithm [26-28] with a “smoothness” value, λ = 10^−5^ (recommended for 64 channels [25]). We refer to the surface-Laplacian transformed EEG signals that represent the macroscopic current sources/sinks on the dura surface under the 60 scalp sites (with the four noise-prone sites excluded from analyses) simply as EEG signals. These EEG-recording and pre-processing procedures were similar to those used in our prior studies [29-31].

### EEG analysis

#### Taking the temporal derivative

We needed to track the peak-alpha frequency and power at each site at a sub-second’s timescale and reasonable spectral resolution. The general 1/f^β^ spectral background in EEG may interfere with the identification of the frequencies and powers at spectral maxima. A commonly employed strategy to circumvent this problem is to compute a Fourier transform over partially overlapping time windows of several seconds or longer, and then fit a power function to each time-windowed Fourier transform to remove the 1/f^β^ component [e.g., 32-34]. However, it is difficult to reliably estimate the 1/f^β^ component on a sub-second’s timescale. Note that β in the 1/f^β^ spectral background varies around 1 (see [35] for a review of the various factors that influence β, and [36] for contributions of the excitatory and inhibitory dynamics to β). Although taking the temporal derivative of EEG (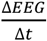 where Δ*t* is the temporal resolution, i.e., 1/512 sec) would completely neutralize the1/f^β^ component only when β = 1, the method has worked well in our prior studies [e.g., 29-31]. Further, as discussed in our prior reports [e.g., 29-31], taking the EEG temporal derivative offers additional advantages. For example, EEG temporal derivatives are drift free. EEG temporal derivatives may be considered a “deeper” measure of neural activity than EEG in the sense that scalp-recorded potentials are generated by the underlying neural currents and taking the EEG temporal derivative macroscopically estimates those currents (as currents in an RC circuit are proportional to the temporal derivative of the corresponding potentials). Further, there is some evidence suggesting that EEG temporal derivatives offer a more effective neural measure than EEG for brain-computer interface use [37].

#### Time-frequency decomposition using Morlet wavelets

To track EEG power spectra at high temporal and spectral resolution, we used a Morlet wavelet-convolution method suitable for time-frequency decomposition of signals containing multiple oscillatory sources of different frequencies (see [25] for a review of different methods for time-frequency decomposition). Each Morlet wavelet is a Gaussian-windowed complex sinusoidal template characterized by its frequency as well as its temporal and spectral widths that limit its temporal and spectral resolutions, respectively. We convolved each EEG waveform (i.e., its temporal derivative) with a set of wavelets tuned to a range of frequencies, yielding a time series of complex values per wavelet frequency. The power and phase of each extracted sinusoidal component at each time point were then given by the modulus squared (power) and the arc tangent of the ratio of the imaginary to the real component (phase). We used a set of wavelets with 160 frequencies, *f*_*w*_’s, ranging between 5 Hz and 15 Hz. The *f*_*w*_’s were logarithmically spaced as neural temporal-frequency tunings tend to be approximately logarithmically scaled [38,39]. The accompanying *n* factor (roughly the number of cycles per wavelet, with the precise definition, *n* = 2*πf · SD*, where *SD* is the wavelet standard deviation) was also logarithmically spaced between 11.7 and 35. This spacing yielded a temporal resolution of *SD* = 370 ms and a spectral resolution of, *FWHM* (*full width at half maximum of wavelet spectrum*) = 1.0 Hz, that were virtually invariant across the range of wavelet frequencies.

We thus obtained the power spectrum in the 5 Hz-15 Hz range as a function of time with the temporal resolution of 370 ms and spectral resolution of 1.0 Hz. To track peak alpha, in each power spectrum (per time point), we identified the curvature maximum that had the highest power, then registered the corresponding frequency and power as the peak-alpha frequency and peak-alpha power. Although we could have simply identified the spectral peak with the highest power, we observed that curvature maxima identified oscillatory frequencies with greater sensitivity and precision than spectral peaks [40].

## Results

We tracked the peak-alpha frequency and power at each site at 370 ms temporal resolution and 1.0 Hz spectral resolution. In our time-averaged data, we replicated the negative posterior-to-anterior gradient of peak-alpha frequency previously reported using MEG [5] in all our behavioral conditions using EEG. The peak-alpha frequency decreased from ∼11 Hz in posterior regions to ∼9 Hz in anterior regions along the midline (Fig 1A-1). We also replicated the typical finding of elevated peak-alpha power in posterior regions when the eyes are closed (Fig 1A-2).

**Fig 1.**
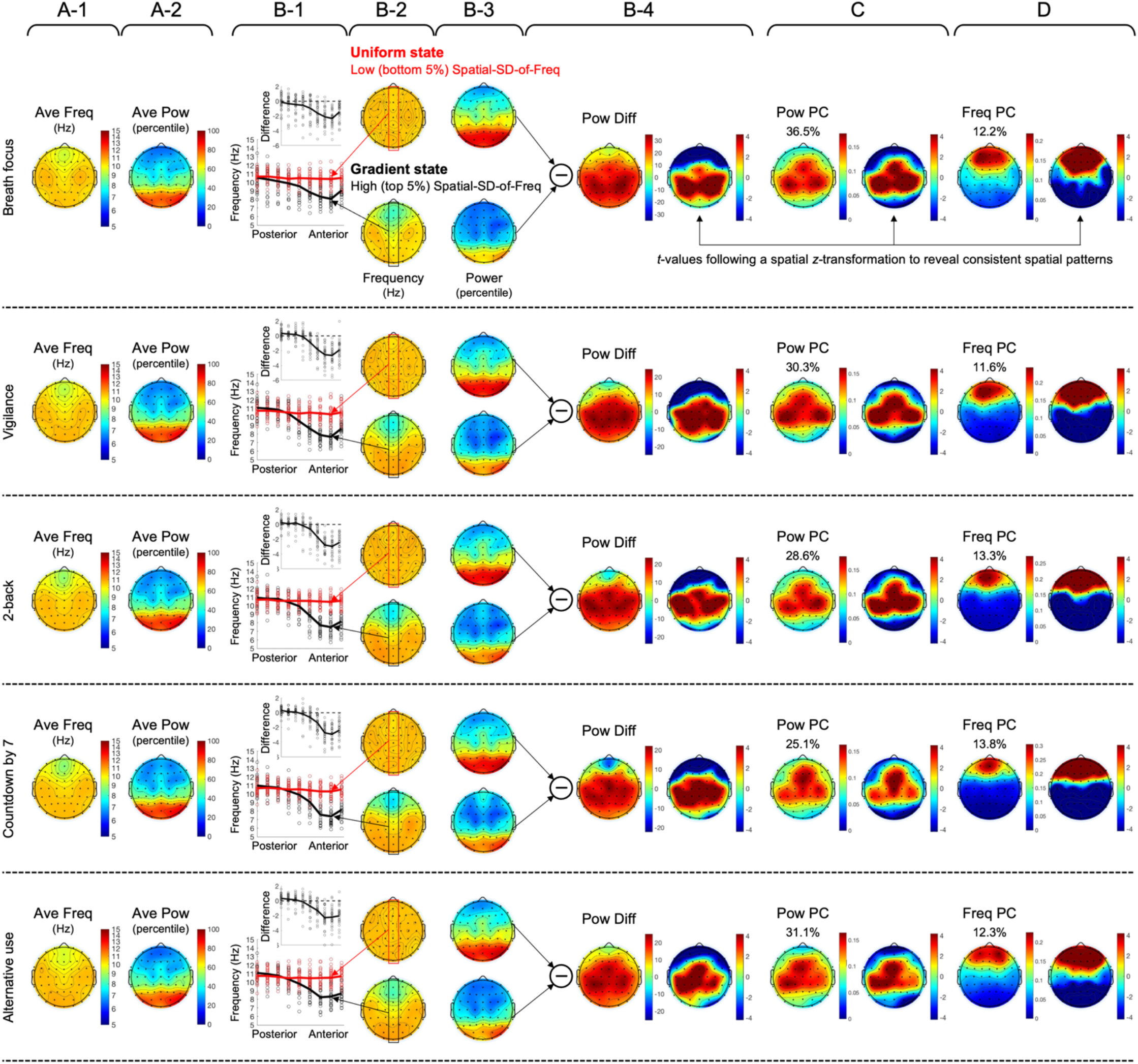
Various spatial maps (columns) of peak-alpha frequency and power for the five behavioral conditions (rows). **A**. The spatial distribution of time-averaged peak-alpha frequency (Hz), *Ave Freq* (**A-1**), and the spatial distribution of time-averaged peak-alpha power (in percentile), *Ave Pow* (**A-2**). Note that Ave Freq diminished from posterior to anterior along the midline (A-1) whereas Ave Pow was elevated in posterior regions (A-2). **B**. The spatial distributions of peak-alpha frequency (Hz) averaged for the periods of low (bottom 5%) spatial standard deviation (**B-2, upper**) and high (top 5%) spatial standard deviation (**B-2, lower**), with the values along the midline axis plotted in **B-1** (the circles representing individual participants) with the upper inset showing the difference between the high- and low-spatial-standard-deviation states. **B-3** shows the spatial distributions of peak-alpha power (in percentile) averaged for the periods of low and high spatial standard deviation of peak-alpha frequency, with their difference, *Pow Diff*, shown in **B-4**, accompanied by the right plot showing *t*-values (following a spatial *z*-transformation) indicating the statistically reliable spatial pattern (|*t*| > 3.75 indicating Bonferroni-corrected statistical significance at *α* = 0.05, 2-tailed). Note that the peak-alpha frequency globally converged at ∼11 Hz in the low-spatial-standard-deviation state, the *uniform state* (B-2, upper, and B-1 red curve), whereas a robust negative posterior-to-anterior gradient emerged in the high-spatial-standard-deviation state, the *gradient state* (B-2, lower, and B-1 black curve). Also note that the peak-alpha power was broadly elevated in central regions in the uniform state relative to the gradient state (B-4). **C**. The spatial loading of the first principal component of the spatiotemporal variations in peak-alpha power, *Pow PC*, with the right plot showing *t*-values (following a spatial *z*-transformation) indicating the statistically reliable spatial pattern (|*t*| > 3.75 indicating Bonferroni-corrected statistical significance at *α* = 0.05, 2-tailed). **D**. The spatial loading of the first principal component of the spatiotemporal variations in peak-alpha frequency, *Freq PC*, with the right plot showing *t*-values (following spatial *z*-transformation) indicating the statistically reliable spatial pattern (|*t*| > 3.75 indicating Bonferroni-corrected statistical significance at *α* = 0.05, 2-tailed). The percentage values shown in C and D indicate the percentages of the variances in peak-alpha power and frequency explained by the corresponding first principal components.

The goal of the current study was to gain insights into how the posterior-to-anterior peak-alpha frequency gradient dynamically formed and dissolved. To this end, we tracked the spatial standard deviation of peak-alpha frequency and examined the spatial distributions of peak-alpha frequency and power when peak-alpha frequency was minimally spatially variable (bottom 5% in spatial standard deviation) and when it was maximally spatially variable (top 5% in spatial standard deviation). It is clear that the instances of low spatial standard deviation corresponded to spatially homogeneous peak-alpha frequency—the *uniform state* (Fig 1B-2, upper)—whereas the instances of high standard deviation corresponded to enhanced posterior-to-anterior gradients—the *gradient state* (Fig 1B-2, lower). In all behavioral conditions, peak-alpha frequency converged at ∼11 Hz along the midline axis in the uniform state (Fig 1B-1, red curve) whereas it decreased from ∼11 Hz (posterior) to 7 Hz-8 Hz (anterior) in the gradient state (Fig 1B-1, black curve). The circles in Fig 1B-1 represent data from individual participants and the upper insets show the difference (the gradient state minus the uniform state).

These results demonstrate that (1) the spatial organization of peak-alpha frequency fluctuates between the uniform and gradient states and (2) the spatial standard deviation of peak-alpha frequency can be used to track the dynamic fluctuations between the uniform and gradient states. We conducted a series of analyses to elucidate the processes that drive the dynamic formations of the uniform and gradient states.

Interestingly, the peak-alpha power was consistently elevated in the uniform state relative to the gradient state (Fig 1B-3 upper vs. lower). Taking the difference showed that the peak-alpha power was broadly elevated in central regions in the uniform state in all behavioral conditions (Fig 1B-4, left). To statistically evaluate the spatial distribution of the central power elevation (in the uniform state), we plotted *t*-values after spatially *z*-transforming power values per participant (to normalize the spatial pattern per participant) (Fig 1B-4, right). Given that |*t*| = 3.34 corresponds to Bonferroni-corrected statistical significance at *α* = 0.05 (two-tailed), the analysis confirmed that the peak-alpha power was broadly elevated in central regions in the uniform state relative to the gradient state (Fig 1B-4, right).

Note that we analyzed peak-alpha power in percentile. As sinusoidal power in human EEG tends to be approximately exponentially distributed [e.g., 29,41], the percentile transformation emphasized power variations in the low to middle ranges. The reason why we chose percentile transformed power for our analyses was that the variations in peak-alpha power were more strongly correlated with the state fluctuations (between the uniform and gradient states) when we used percentile (rather than raw) power values. Specifically, we derived a univariate time series of spatiotemporal peak-alpha power variations most likely associated with the state fluctuations. We did so by computing a dot-product (in the spatial dimension) between the spatial pattern of peak-alpha power difference between the uniform and gradient states—the Pow Diff template (Fig 1B-4 but computed separately for each participant)—and the spatiotemporal variations in peak-alpha power. In other words, we multiplied the site vector (row vector) representing the Pow Diff template by the timepoint-by-site matrix of spatiotemporal peak-alpha power (per condition per participant), yielding the Pow Diff time series likely associated with the state fluctuations. We computed the Pow Diff time series using both raw and percentile power values and examined their correlations with the state fluctuations (i.e., with the temporal variations in the spatial standard deviation of peak-alpha frequency). The correlation was consistently stronger (i.e., more negative) using percentile power than using raw power, indicated by the difference, *r*_percentile_ – *r*_raw_, being consistently negative for nearly all participants in all behavioral conditions (Fig 2). This suggests that the state fluctuations between the uniform and gradient states are primarily associated with peak-alpha power variations in the low to middle range (emphasized by the percentile transformation) in central regions (Fig 1B-4).

**Fig 2.**
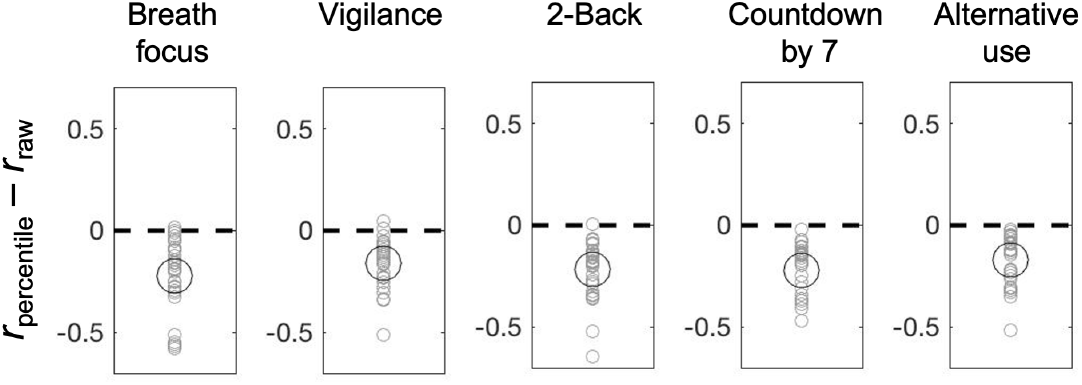
Correlation between the spatial standard deviation of peak-alpha frequency and peak-alpha power in central regions computed with percentile versus raw power. For each participant for each behavioral condition, we computed the difference in Pearson’s *r* (Fisher-*z* transformed) between the value computed with percentile power and that computed with raw power, *r*_percentile_ minus *r*_raw_. Note that the negative correlation (higher peak-alpha power in central regions being associated with lower spatial standard deviation of peak-alpha frequency, i.e., the uniform state) was consistently stronger using percentile power than using raw power for nearly all participants (gray circles) in all behavioral conditions. The large black circles represent the means.

As expected, the spatial standard deviation of peak-alpha frequency was consistently negatively correlated with the Pow Diff time series indicating that higher peak-alpha power in central regions were consistently associated with lower spatial standard deviation of peak-alpha frequency (or with the uniform state) for all participants in all behavioral conditions (Fig 3A, leftmost gray circles). Because the Pow Diff template was derived as the spatial pattern of peak-alpha power difference between the uniform and gradient states, it served as a benchmark to evaluate the contributions of spatially coordinated peak-alpha power variations to the state fluctuations.

**Fig 3.**
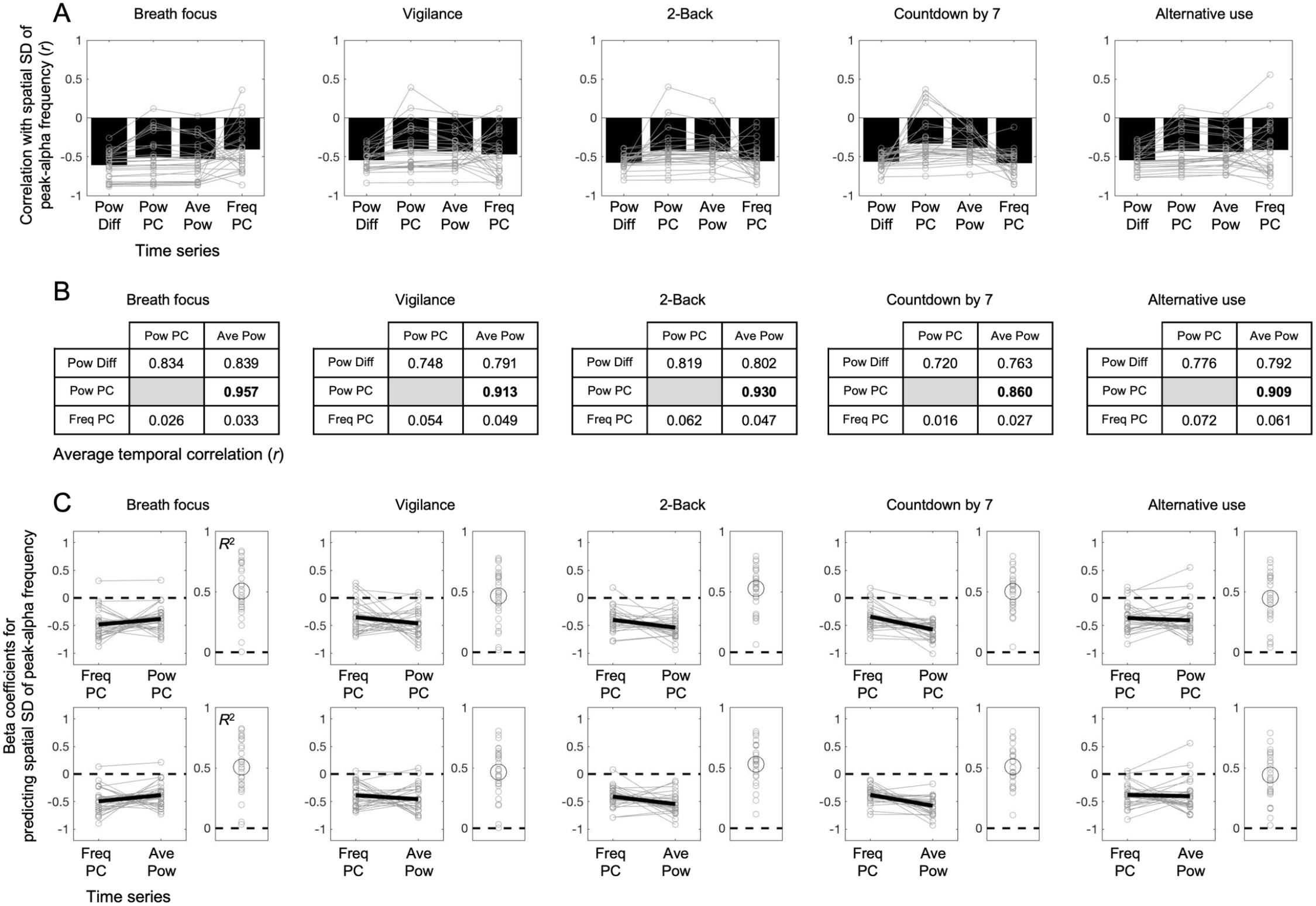
Correlations between the spatial standard deviation (SD) of peak-alpha frequency and various spatial components of peak-alpha power and frequency for the five behavioral conditions (columns). **A**. Correlations (Pearson’s *r*) between the spatial SD of peak-alpha frequency and the Pow Diff time series (leftmost), the Pow PC time series (mid-left), Ave Pow time series (mid-right), and the Freq PC time series (rightmost) (see text). The gray circles connected with lines represent individual participants and the black bars represent the means. Note that all four time-series correlated negatively with the spatial SD of peak-alpha frequency. Also note that the correlations with the Pow Diff, Pow PC, and Ave Pow time series showed nearly identical individual differences (the gray lines being approximately parallel) suggesting that these variables are redundant, whereas the individual differences were not preserved for the correlations with the Freq PC time series suggesting that this variable is distinct. **B**. Average pairwise correlations among the Pow Diff, Pow PC, Freq PC, and Ave Pow time series. Note that the correlations were high among the Pow Diff, Pow PC, and Ave Pow time series, and especially high between the Pow PC and Ave Pow time series (bolded). In contrast, the Freq PC time series minimally correlated with either the Pow PC time series or the Ave Pow time series. **C**. Predicting the spatial SD of peak-alpha frequency with the Freq PC and Pow PC time series (upper left panels) or with the Freq PC and Ave Pow time series (lower left panels). Beta coefficients are shown with the gray lines representing individual participants and the thick black line representing the mean. The proportion of temporal variance of the spatial SD of peak-alpha frequency accounted for by each pair of variables is shown as *R*^2^ (right panels) with the gray circles representing individual participants and the large black circle representing the mean. Note that nearly all participants yielded negative beta coefficients for both variables of each pair with each pair accounting for a substantial proportion (∼50% on average) of the temporal variance of the spatial SD of peak-alpha frequency.

To identify the contributing spatially coordinated power variations, we evaluated two spatial templates, (1) the spatial loading of the first principal component of the spatiotemporal variations in peak-alpha power—the *Pow PC* template (Fig 1C but computed separately for each participant)—and (2) the spatial pattern of time-averaged peak-alpha power—the *Ave Pow* template (Fig 1A-2 but computed separately for each participant). The Ave Pow template reflected peak-alpha power variations in posterior regions where peak-alpha power was generally elevated (Fig 1A-2). The Pow PC template (i.e., the first principal component) accounted for 25%-36% of the spatiotemporal variance in peak-alpha power and showed a central spatial distribution similar to the benchmark Pow Diff template (compare Fig 1C with Fig 1B-4).

We used the Pow PC and Ave Pow templates to convert the spatiotemporal peak-alpha power variations into univariate time series (by computing dot-products in the spatial dimension as described above). We refer to the Pow PC-template convolved and Ave Pow-template convolved time series as the Pow PC time series and Ave Pow time series, respectively. Both time series negatively correlated with the state fluctuations to a similar degree to the benchmark Pow Diff time series for most participants in all behavioral conditions (Fig 3A, largely parallel gray lines through the Pow Diff, Pow PC, and Ave Pow time series). This result is consistent with the interpretation that the state fluctuations were driven by spatially coordinated peak-alpha power variations reflected in the Pow PC and Ave Pow templates. Indeed, the Pow Diff, Pow PC, and Ave Pow time series were highly correlated with one another (Fig 3B). The particularly strong correlation between the Pow PC and Ave Pow time series (*r* = 0.860 to 0.975 in the five behavioral conditions; bolded in Fig 3B) suggest that the coordinated power variations in central and posterior regions equivalently contributed to the state fluctuations.

Given that peak-alpha power variations in central (detected by the Pow PC template) and posterior (detected by the Ave Pow template) regions were equivalently correlated with the state fluctuations, we examined the cross correlation between the two variables to see whether one region tended to drive the other region. For each participant (and each behavioral condition), we computed the time shift of the Ave Pow time series relative to the Pow PC time series at their cross-correlation peak. A negative value indicated that the Pow PC time series varied ahead of the Ave Pow time series whereas a positive value indicated that the Ave Pow time series varied ahead of the Pow PC time series. Although there were individual differences, the distribution of the time shift was in the positive direction for most participants in all behavioral conditions (Fig 4), suggesting that peak-alpha power variations in posterior regions (detected by the Ave Pow template) drive peak-alpha power variations in central regions (detected by the Pow PC template).

**Fig 4.**
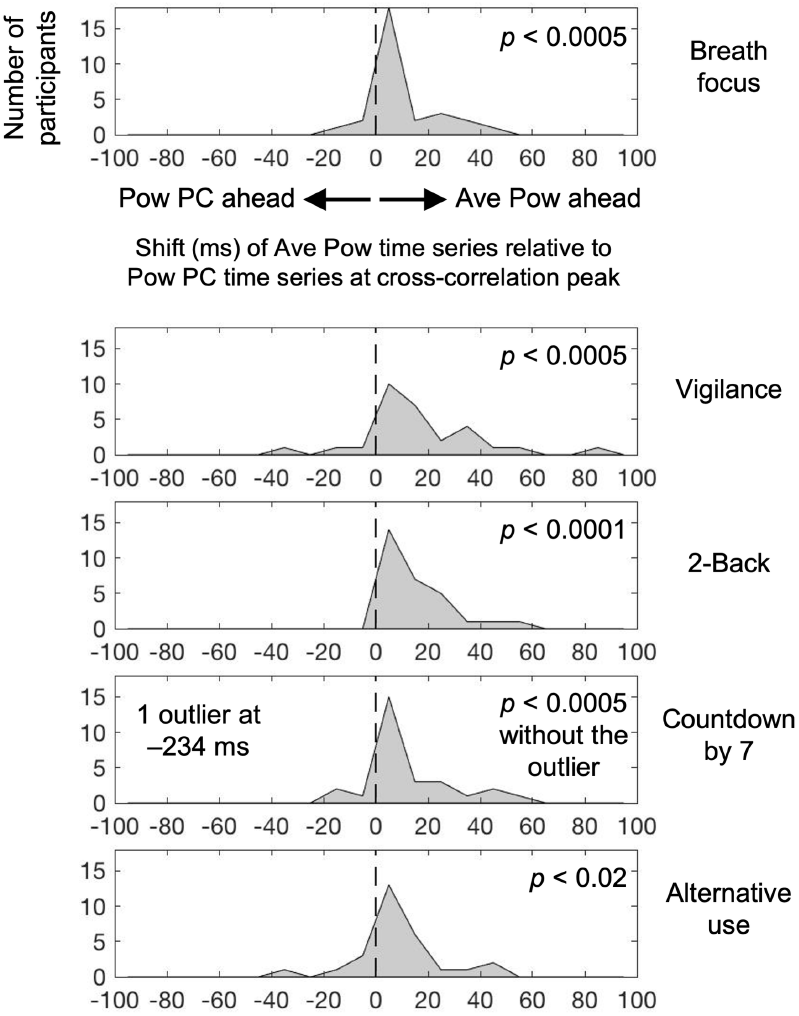
Distribution (across participants) of the temporal peak of cross correlation between the Pow PC and Ave Pow time series for the five behavioral conditions. Note that the distribution is generally positively shifted in all behavioral conditions, indicating that the posteriorly localized Ave Pow time series (see Fig 1A-2) varied ahead of the centrally localized Pow PC time series (see Fig 1C). This suggests that coordinated variations in peak-alpha power in posterior regions drive those in central regions.

Our analysis so far suggests that coordinated posterior variations in peak-alpha power that spread to central regions drive the state fluctuations between the uniform and gradient states. Interestingly, we also identified coordinated anterior variations in peak-alpha frequency, reflected in the first principal component, that accounted for 11.6% to 13.8% of the spatiotemporal variance in peak-alpha frequency. The spatial loading of this principal component, Freq PC (Fig 1D), showed that the coordinated peak-alpha frequency variations were localized in anterior regions similar to the anterior dip in peak-alpha frequency characteristic of the gradient state (see lower plots in Fig 1B-2). This observation suggested that the coordinated anterior frequency variations also drive the state fluctuations either independently or in coordination with the coordinated posterior-to-central power variations. Indeed, spatiotemporal variations in peak-alpha frequency convolved with the Freq PC template (Fig 1D but computed separately for each participant), which we refer to as the Freq PC time series, consistently negatively correlated with the state fluctuations for most participants in all behavioral conditions (Fig 3A for Freq PC).

The gray lines (representing individual participants) in Fig 3A are overall parallel for the Pow Diff, Pow PC, and Ave Pow time series, suggesting that the peak-alpha power variations convolved with these templates similarly negatively correlated with the state variations. In contrast, the gray lines substantially crossed in relation to the Freq PC time series, suggesting that the peak-alpha frequency variations convolved with the Freq PC template accounted for different dynamics of the state fluctuations. Indeed, the Freq PC time series were minimally correlated with either the Pow PC times series or the Ave Pow time series (Fig 3B, bottom rows). Thus, the state fluctuations between the uniform and gradient states appear to be driven by two separate processes, (1) coordinated variations in peak-alpha power spreading from posterior to central regions and (2) coordinated variations in peak-alpha frequency in anterior regions. To confirm this observation, we ran linear multiple regression models predicting the state fluctuations using the Freq PC time series in combination with either the Pow PC or the Ave Pow time series.

Most participants in all behavioral conditions yielded negative beta coefficients for both the Freq PC and Pow PC time series (Fig 3C, gray circles in upper left panels) as well as for both the Freq PC and Ave Pow time series (Fig 3C, gray circles in lower left panels). Further, whereas the Pow PC, Ave Pow, and Freq PC time series each accounted for ∼25% of the variance (on average) in the state fluctuations (Fig 3A, showing *r* ∼ 0.5 on average), the Freq PC-Pow PC pair as well as the Freq PC-Ave Pow pair accounted for ∼50% of the variance (on average) in the multiple regression models (Fig 3C, right panels). These results confirm that coordinated variations in peak-alpha power spreading from posterior to central regions and coordinated variations in peak-alpha frequency in anterior regions relatively independently contribute to driving the spatial organization of peak-alpha frequency between the uniform and gradient states.

## Discussion

Prior research has demonstrated a generally negative posterior-to-anterior gradient of peak-alpha frequency, slowing from posterior to anterior regions [5,15], and associated with increases in cortical thickness [5]. Increases in cortical thickness have been linked to increases in connectivity [6] and the ratio of feedback to feedforward connections [7]. Further, regions with increased connectivity tend to show slower fMRI BOLD fluctuations [8,9], the time scale of intrinsic fluctuations in spiking activity tends to increase from sensory areas to association areas in primate cortex [10], and the cortical gradient of connectivity is similar to that of temporal integration, generally increasing from posterior to anterior regions [11,12]. Taken together, these relations suggest that the negative posterior-to-anterior gradient of peak-alpha frequency reflects the cortical hierarchy in temporal integration [5,13]. Given that this gradient may be accompanied by traveling waves from posterior to anterior regions [14,15], the gradient may facilitate information influx into anterior regions in addition to increasing temporal integration toward anterior regions. We consistently replicated the general negative posterior-to-anterior gradient previously reported in MEG [5], in scalp recorded EEG while participants engaged in a variety of behavioral tasks with their eyes closed, breath focus, vigilance, working memory, mental arithmetic, and generative thinking.

A computational model of Macaque brain showed that the posterior-to-anterior hierarchy in temporal integration could arise from a combination of heterogeneity in excitatory connection strengths across regions and specific profiles of long-range connectivity [13]. The model further showed that this hierarchy could be dynamically adjusted by differentially gating long-range inputs, raising the possibility that the negative posterior-to-anterior gradient of peak-alpha frequency may be dynamically controlled (also suggested by [5]).

The unique contribution of the current study is the demonstration that the spatial organization of peak-alpha frequency fluctuates between the gradient state where peak-alpha frequency forms a robust negative gradient, diminishing from ∼11 Hz in posterior regions to ∼8 Hz in anterior regions, and the uniform state where it globally converges at ∼11 Hz. Global frequency convergence in alpha-band oscillations may facilitate global neural communication by allowing phase locking [e.g., 42]. Thus, alternations between the uniform and gradient states may reflect alternations between promoting global communication and imposing a posterior-to-anterior hierarchy in temporal integration and information flow.

Importantly, our results suggest that the fluctuations between the uniform and gradient states are controlled by two distinct (uncorrelated) processes. One process, underlying the first principal component of the spatiotemporal variations in peak-alpha frequency, appears to directly control peak-alpha frequency in anterior regions. This anterior process may control alpha-oscillation frequencies by adjusting temporal integration (e.g., longer integration promoting slower oscillations) through appropriately adjusting the weights of relevant long-range inputs [e.g., 13]. The other process, underlying the first principal component of the spatiotemporal variations in peak-alpha power, appears to indirectly control peak-alpha frequency through facilitating synchronization. Specifically, this process appears to initiate a wave of oscillatory entrainment in the upper alpha band originating in posterior regions that spreads to central regions to promote global entrainment in the upper alpha band.

In conclusion, we replicated the negative posterior-to-anterior gradient in peak-alpha frequency observed in MEG [5] in time-averaged EEG data while participants engaged in a variety of cognitive tasks, consistent with the idea that the peak-alpha-frequency gradient reflects the general posterior-to-anterior hierarchy in temporal integration [e.g., 5,12,13]. The current results suggest that the spatial pattern of peak-alpha frequency dynamically fluctuates between the gradient state, potentially facilitating information influx and temporal integration toward anterior regions, and the uniform state, potentially facilitating global communication. Importantly, the state fluctuations are controlled by at least two distinct mechanisms, an anterior mechanism that directly adjusts peak-alpha frequencies and a posterior mechanism that indirectly adjusts them by increasing or decreasing global entrainment in the upper alpha band. Future research is required to specify the neural computations underlying these anterior and posterior mechanisms as well as their functional roles.

## Notes

https://doi.org/10.21985/n2-d0bt-a448

### Competing Interest Statement

The authors have declared no competing interest.

